# The conformation of the nSrc specificity-determining loops in the Src and Abl SH3 domains are modulated by a “WX” conserved sequence motif

**DOI:** 10.1101/2024.08.26.609727

**Authors:** Frederick Longshore-Neate, Caroline Ceravolo, Cole Masuga, Elise F. Tahti, Jadon M. Blount, Sarah N. Smith, Jeanine F. Amacher

**Affiliations:** Department of Chemistry, Western Washington University, Bellingham, Washington, 98225, United States

**Keywords:** SH3 domains, protein-protein interactions, protein-peptide interactions, short linear motifs, specificity, signal transduction

## Abstract

Cellular signaling networks are modulated by multiple protein-protein interaction domains that coordinate extracellular inputs and processes to regulate cellular processes. Several of these domains recognize short linear motifs, or SLiMs, which are often highly conserved and are closely regulated. One such domain, the Src homology 3 (SH3) domain, typically recognizes proline-rich SLiMs and is one of the most abundant SLiM-binding domains in the human proteome. These domains are often described as quite *versatile*, and indeed, SH3 domains can bind ligands in opposite orientations dependent on target sequence. Furthermore, recent work has identified diverse modes of binding for SH3 domains and a wide variety of sequence motifs that are recognized by various domains. Specificity is often attributed to the RT and nSrc loops near the peptide-binding cleft in this domain family, particularly for Class I binding, which is defined as RT and nSrc loop interactions with the N-terminus of the ligand. Here, we used the Src and Abl SH3 domains as a model to further investigate the role of the RT and nSrc loops in SH3 specificity. We created chimeric domains with the loop sequences swapped between these SH3 domains, and used fluorescence anisotropy assays to test how relative binding affinities were affected for Src SH3- and Abl SH3-specific ligands. We also used Alphafold – Multimer to model our SH3:peptide complexes. We identified a position that contributes to the nSrc loop conformation, the amino acid immediately C-terminal to a highly conserved Trp that creates a hydrophobic pocket critical for SH3 ligand recognition. We defined this as the WX motif, where X = Trp for Src and Cys for Abl. The importance of this position for orienting the ligand is supported by analyses of previously deposited SH3 structures, multiple sequence alignment of SH3 domains in the human proteome, and our biochemical data of mutant Src and Abl SH3 domains. Overall, our work uses experimental approaches and structural modeling to better understand SH3 specificity determinants.

## Introduction

Short linear motif (SLiM) or peptide binding is a critical component of signal transduction pathways, often with several SLiM-binding domains in multiple proteins modulating the activity and regulation of the signaling cascade (*1*). One such domain, the Src-homology 3 (SH3) domain, was first described in 1988 as a domain located on the proto-oncogene Src tyrosine kinase (*2*, *3*). It is now recognized that SH3 domains exist in all kingdoms of life and viruses, and there are over 300 SH3 domains in 200 proteins in the human proteome (*4*, *5*). Although several noncanonical exceptions exist, SH3 domains are generally characterized as binding to proline-rich sequences, specifically containing a PXXP motif (where X = any amino acid), with affinities in the low-to-mid micromolar range (*5*, *6*). Many SH3 target sequences adopt a type II polyproline helix (PPII) structure, presenting a hydrophobic surface to which the SH3 domain recognizes and binds (*7*, *8*).

SH3 domains share a general conserved fold, despite displaying a relatively large degree of plasticity in ligand binding. SH3 domains are typically about 60 amino acids in length, with a β-barrel fold, consisting of approximately 5 β--strands and a 310 helix (*4*, *10*) (**Figure 1**). Specificity in SH3 domain binding is determined by the n-Src, RT, and β4-α310 loops (*4*). Interestingly, the same SH3 domain can bind ligands in opposite orientations, depending on whether it corresponds to a Class I/“plus” (consensus sequence: RXLPPXP, where X=any amino acid) or Class II/“minus” (XPPLPXR) target sequence (*7*, *11–13*). In the Class I orientation, N-terminal residues of the ligand interact directly with the RT and n-Src loops; in the Class II orientation, these regions interact with C-terminal residues of the ligand (*7*, *11*). Thus, the same SH3 domain can bind ligands N- to C-terminal or vice versa depending on the target sequence.

**Figure 1.**
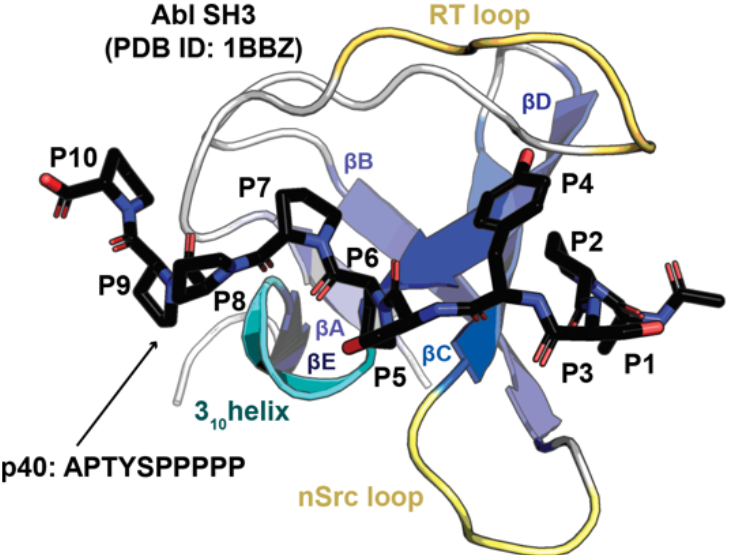
Structure of an SH3 domain. The Abl SH3 structure is shown as a cartoon bound to the high affinity p40 (APTYSPPPPP) peptide, which is shown in stick representation and colored by atom (C=black, O=red, N=blue), PDB ID: 1BBZ (9). Conserved structural elements are highlighted and labeled, including the βA-βE strands, 310 helix, and RT and nSrc loops. Peptide positions, P1-P10, are labeled.

The *Src module*, consisting of an SH3 domain, SH2 domain, and tyrosine kinase, is shared amongst several families of cytoplasmic tyrosine kinases (*14*). This includes Abl, a Src-related tyrosine kinase, which regulates actin (*15*). The kinase domains of Src and Abl are 46% identical, but these proteins are known to be differentially regulated (*16–19*). Furthermore, therapeutics that target the BCR-Abl fusion protein, which is the underlying cause of disease in approximately 95% of chronic myelogenous leukemia (CML) cases, e.g., imatinib, bind and inhibit Abl kinase, but do not bind Src kinase (*16*). There are also specificity differences in the SH3 domains of these proteins, which are 43% identical over 58 residues. A previous study of SH3 domain specificity investigated an initial ligand, derived from the 3BP1 protein (sequence: RAPTMPPPLPP), which bound the Abl and Fyn (a Src family kinase) SH3 domains with similar 30 μM affinity (*6*, *9*). This sequence was then used as a template to design a selective and high affinity Abl SH3 ligand, termed p40 (*6*, *9*). The Class I p40 ligand (APTYSPPPPP) bound the Abl and Fyn SH3 domains with 0.4 μM and 470 μM affinity, respectively (*6*, *9*).

Here, we aimed to better understand this result by expanding this investigation to Abl and Src SH3 specificity using chimeric proteins, Abl_Src_ and Src_Abl_, which swap the RT and n-Src loop sequences from the other domain. We calculated binding affinities of these chimeric domains with the Class I p40 ligand, as well as a Class I Src-specific target sequence, LASRPLPLLP (*20*). We found that while the WT Src and Abl SH3 domains are specific for their ligands, the Src_Abl_ chimeric protein revealed relatively weak but similar binding to both peptide sequences. To visualize SH3:ligand interactions, we used Alphafold – Multimer to model the complexes and analyze potential binding interfaces. We also tested single mutations of a WW (for Src SH3) and WC (for Abl SH3) sequence motif at the C-terminus of the n-Src loop in the wild-type and chimeric proteins, showing that these can modulate ligand affinity several fold. These positions in Src SH3, the amino acids W121 and W122, have previously been shown to be important for establishing a hydrophobic binding specificity pocket and modulating lipid binding (*21*, *22*). Here, we argue that the W122 position additionally controls the conformation of the nSrc loop, which affects ligand positioning and specificity. Taken together, our studies provide further insight into the role of these specificity-determining loops for the Src and Abl SH3 domains.

## Results

### Differing specificities of Src and Abl SH3 domains

To investigate the effect of RT and n-Src loop variation on the Src and Abl SH3 domains, we first wanted to confirm that these domains contain differing ligand specificities. We recombinantly expressed and purified the Src and Abl SH3 domains as SUMO-tagged fusion proteins, as described in the Materials and Methods. All sequences used in this work are in the **Supporting Information**. We then used fluorescence anisotropy experiments, as described previously and in the Materials and Methods, to test the binding affinities of each domain with two fluorescein (*F\**)-tagged peptides, one corresponding to the p40 sequence, *F\**-Ahx-APTYSPPPPP, termed *F\**-p40, and to a Src SH3 identified ligand, *F\**-Ahx-LASRPLPLLP, termed *F\**-PLLP (**Figure 2**, **Table 1**) (*6*, *9*, *20*).

**Figure 2.**
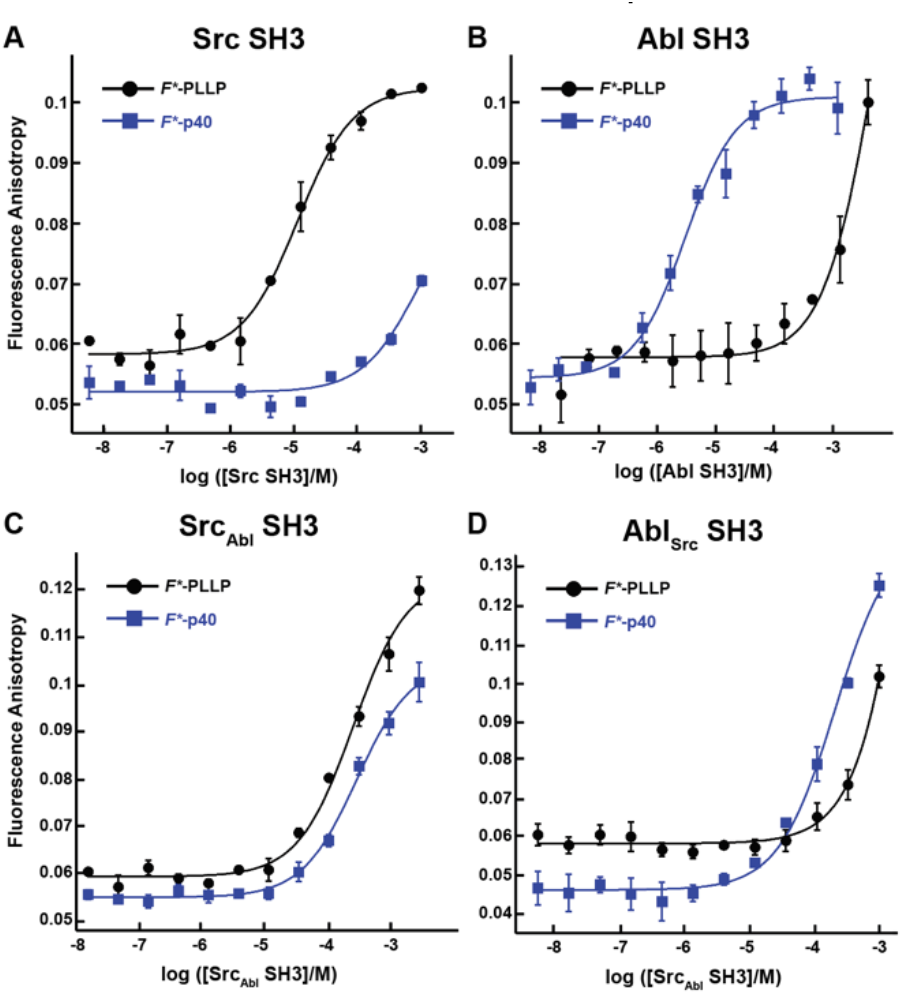
Binding affinities for F*-PLLP and p40 with Src, Abl, Src_Abl_, and Abl_Src_ SH3 domains. The binding curves for the Src (**A**), Abl (**B**), Src_Abl_ (**C**), and Abl_Src_ (**D**) SH3 domains with the F*-PLLP (LASRPLPLLP) and F*-p40 (APTYSPPPPP) peptides Averaged values and standard deviations are shown for each and calculated binding affinities are labeled and are in **Table 1**. Because of differing protein concentrations, the data in these graphs may not include all replicates used for binding affinity measurements.

**Table 1.**
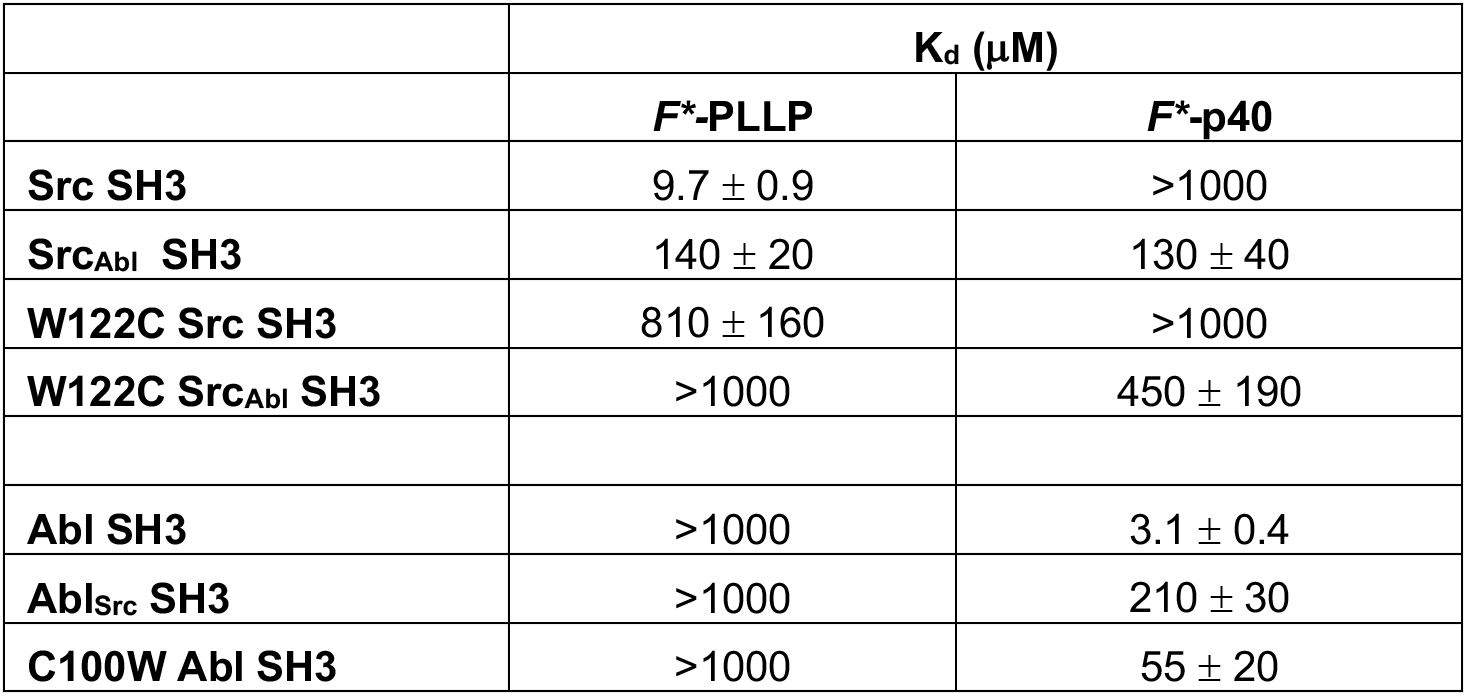
Relative binding affinities using fluorescence anisotropy for Src- and Abl-derived SH3 domains.

Consistent with previous work, our results indicated that while Src SH3 bound *F\**-PLLP with relatively high affinity, *K*D = 9.7 ± 0.9 μM, it did not bind *F\**-p40, here defined as *K*D > 1000 μM (**Figure 2A**, **Table 1**). In contrast, Abl SH3 bound *F\**-p40 with *K*D = 3.1 ± 0.4 μM, and *F\**-PLLP with *K*D > 1000 μM (**Figure 2B**, **Table 1**). Notably, our Abl SH3 affinity for *F\**- p40 differs by an order of magnitude as compared to the previously published value, 0.4 ± 0.1 μM (for reference, in this paper Fyn SH3 bound with *K*D = 472 ± 55 μM) (*6*, *9*); however, we attribute these differences to the experimental methods used, variations in sequence, i.e., our peptides also include N-terminal fluorescein and Ahx linkers, and the use of Fyn SH3 as opposed to Src SH3, whose SH3 domains are 77% identical by sequence. Despite variation in absolute values, our results also indicated a difference of approximately three orders of magnitude for Abl SH3 binding to *F*-*p40 as compared to Src/Fyn SH3, indicating internal consistency with the previous results. Taken together, our results confirmed differing ligand specificities for the Abl and Src SH3 domains.

### Chimeric SH3 domains confer differences in ligand specificities

Previous reports indicated that the RT and nSrc loops of SH3 domains drive ligand specificity (*4*). Therefore, we wanted to test chimeric proteins, with the RT (sequence: ^96^ESRTET^101^) and nSrc (sequence: ^117^TEGD^120^) loops of Src swapped for the RT (sequence: ^73^VASGDN^78^) and nSrc (^95^HNGE^98^) loops of Abl, and vice versa in each SH3 domain. We termed the resulting chimeric proteins, Src_Abl_ and Abl_Src_ SH3, where the subscript indicates the identity of the RT and nSrc loops. These proteins were also recombinantly expressed and purified as SUMO fusions and binding affinities determined using fluorescence anisotropy experiments with our *F\**-PLLP and *F\**-p40 peptides, as described in the Materials and Methods (**Figure 2**, **Table 1**).

In our Src_Abl_ SH3 chimera, binding affinity for the Src-specific *F\**-PLLP peptide was reduced ∼14- fold, to *K*D = 140 ± 20 μM, as compared to the wild-type Src SH3 domain (**Figure 2A**, **Table 1**). Interestingly, Src_Abl_ SH3 bound the Abl-specific *F\**-p40 peptide with similar affinity, *K*D = 130 ± 40 μM, indicating that specificity was indeed removed by this substitution (**Figure 2C**, **Table 1**). In contrast, while binding to the Abl-specific *F\**-p40 peptide was reduced by > 60-fold for Abl_Src_ SH3, *K*D = 210 ± 30 μM, this chimeric protein continued to show undetectable binding to the Src-specific *F\**-PLLP peptide, *K*D > 1000 μM (**Figure 2D**, **Table 1**).

### Structural analyses of SH3-ligand models

To better understand our biochemical assay results, we generated structural models of our SH3-ligand complexes using Alphafold – Multimer on the ColabFold server (*23*, *24*). We created models of wild-type Src and Abl SH3 domains with each peptide, PLLP and p40, as well as our chimeric Src_Abl_ and Abl_Src_ SH3 proteins with each peptide (**Figures 3A-B**). Overall, the 5 models generated for each complex were internally consistent, with alignments of < 0.18 Å in all cases (**Figures 3A-B**). However, there were notable exceptions, which included steric clash based on the PyMOL library for several of the peptides with SH3 domains. In addition, two of the models for Abl_Src_ SH3 with the PLLP peptide were modelled in the opposite Class II orientation, again, defined as the C-terminus of the peptide interacting with the RT and nSrc loops (**Figure 3B**) (*7*, *11*, *13*). Therefore, our analyses will be based on models which include the Class I binding orientation and no steric clash.

**Figure 3.**
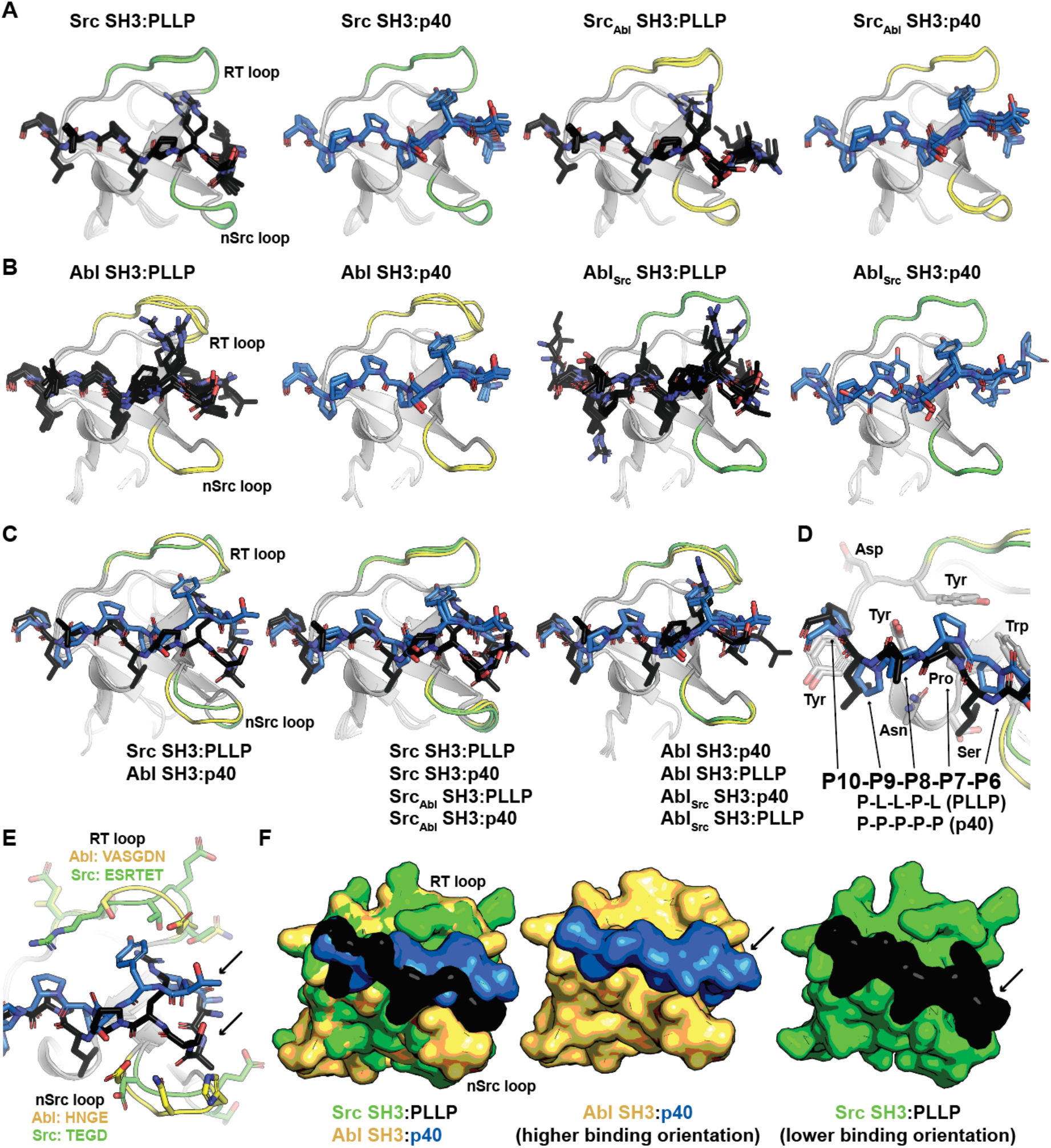
Alphafold – Multimer (ColabFold) models of Src, Abl, Src_Abl_, and Abl_Src_ SH3 domains with F*- PLLP and p40 ligands. (**A-B**) Alphafold – Multimer results for Src and Src_Abl_ (**A**) or Abl and Abl_Src_ (**B**) SH3 domains with the PLLP (black sticks, colored by atom) or p40 (blue sticks, colored by atom) ligands. In all cases, the results for each complex align with RMSD values of < 1.75 Å, with many even closer (e.g., Abl SH3:PLLP aligns with RMSD < 1.3 Å). SH3 domains are shown in cartoon representation with the RT and nSrc loops colored based on the origin sequence, green for Src SH3 and yellow for Abl SH3. The PLLP and p40 peptides are shown in stick representation and colored by atom (C = black (PLLP) or blue (p40)). (**C**) Although the Src and Abl RT and nSrc loops are in distinct conformations (left figure), alignment of the chimeric SH3 domains with their respective wild- type counterparts indicates that the loop conformations largely match the wild-type scaffold. Structures are rendered and colored as in (**A-B**). (**D**) SH3 residues in Src and Abl that interact with the P6-P10 ligand positions are identical. SH3 domains are in cartoon representation with side chain sticks for interacting residues (C = gray). Peptide positions are labeled. (**E**) Differences in the RT and nSrc loops (as shown in **C**) modulate ligand binding, such that peptides bind Abl in a “higher” orientation and Src in a “lower” orientation. In the left figure, the SH3 domains are shown in cartoon, with side chain atoms of the RT and nSrc loops shown as sticks (C = green (Src) or yellow (Abl)). The peptides are rendered as in (**A-D**). (**F**) Surface representation of (**E**). Here, arrows indicate the differing peptide orientations for Src (green), Abl (yellow), PLLP (black), and p40 (blue). In all stick representation images, O = blue, N = red.

While we observed loop conformation differences between the Abl and Src SH3 domains, our Src_Abl_ and Abl_Src_ chimeric protein loops were consistent with the parent/wild-type protein, e.g., the Abl_Src_ loops are in an Abl SH3 conformation (**Figure 3C**). The PXXP motifs at the C-terminus of each peptide (PLLP and PPPP, respectively, and referred to as positions 7-10, or P7-P10) interact largely with the βA-βB loop and side-chains in the 310 helix in a consistent manner between the two peptide sequences and other SH3 domains (*9*). All PxxP-interacting residues are conserved between Abl and Src SH3 (**Figure 3D**). At the N-terminal end of the peptide, positions 1-6 (or P1-P6), differences in the conformation of the nSrc loops revealed differing peptide conformations (**Figures 3E-F**) and an upward translation of the peptide-binding cleft for Abl SH3 as compared to Src SH3 (**Figures 3E-F**).

Analyses of the structures in combination with our binding affinity results suggested that the orientation of the Abl nSrc loop was incompatible with binding to the N-terminal P1-P6 LASRPL sequence of the PLLP peptide. We hypothesized this is due to the lowered translation of the peptide required to allow for an electrostatic interaction between the P4 Arg and D102 Src SH3, using full-length Src numbering (**Figure 4A**). This interaction is a well-studied characteristic of Src SH3 specificity (*12*, *13*, *25*). We will refer to this as the lower “Src-binding” peptide conformation. The D102 amino acid immediately follows the RT loop. In Abl, this position corresponds to T79. The equivalent peptide translation would result in steric clash with the nSrc loop in the elevated “Abl-binding” orientation, as the main chain atoms of the peptide are translated ∼2 Å between the two peptide conformations (**Figure 4B**). There is an Arg residue in the RT loop of Src, R98, which may further facilitate this lowered conformation to avoid steric clash with the P7 Pro in each peptide (**Figure 4C**). However, in the Abl RT loop, this equivalent position is S75, suggesting that the elevated “Abl-binding-like” peptide conformation is accessible to the Src_Abl_ SH3 domain; thus, providing rationale as to why Src_Abl_ SH3 bound both the *F*-*PLLP and *F*-*p40 peptides with relatively equivalent affinities (**Figure 4D**, **Table 1**).

**Figure 4.**
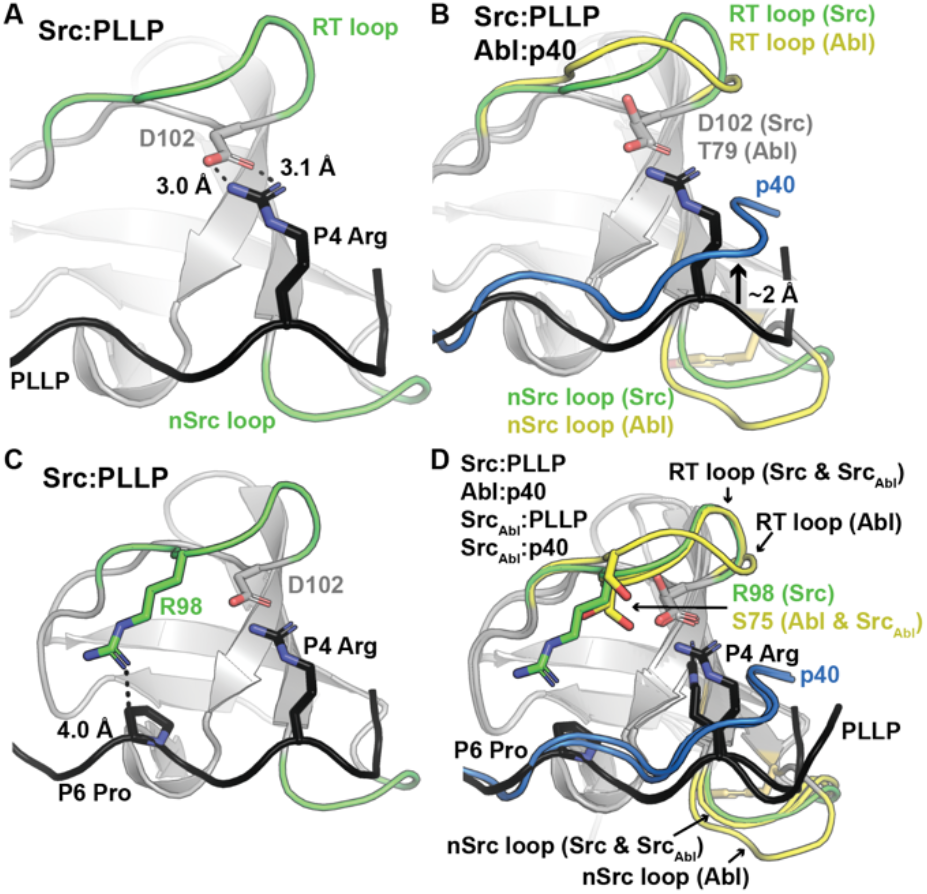
Ligand interactions with the RT and nSrc loops of Src and Abl SH3 domains. In all, SH3 domains and ligands are shown in cartoon representation, with loops colored by original sequence (green for Src and yellow for Abl), even in the chimeric proteins, as labeled. The ligands are shown as black for PLLP and blue for p40. Relevant side chains are shown as sticks and colored by atom (C=as above, N=blue, O=red). (**A**) The P4 Arg in the ligand interacts with D102 in Src SH3. Distances shown as black dashed lines and labeled. (**B**) The p40 ligand bound to Abl SH3 is translated ∼2 Å in the ligand binding pocket. (**C**) The lowered peptide-binding position in Src SH3 is further determined by the presence of R98 in the RT loop, which dictates positioning of the P6 Pro of the ligand. (**D**) Although the RT loop of Src_Abl_ SH3 is in a similar conformation as wild-type Src SH3, R98 (Src) is replaced by S75 (Abl), allowing the p40 peptide to bind Src_Abl_ in the elevated peptide-binding conformation. The nSrc loop of Src_Abl_ is also in a Src-like conformation.

### The WX sequence motif at the base of the nSrc loop influences nSrc loop orientation

Structural analyses of our Alphafold-generated models also suggested that an intra-SH3 interaction on either side of the nSrc loop modulates the respective loop conformations. Specifically, an aromatic Trp (W122) C-terminal to the nSrc loop is counterbalanced by an aromatic Tyr (Y93) N-terminal to the nSrc loop in Abl SH3. We predicted these residues and their interaction with an amino acid on the other side of the loop (N116 in Src SH3 and C100 in Abl SH3), either with side chain or main chain atoms, modulates the differing positions of the nSrc loop we observed (**Figure 5A**). Furthermore, we reasoned that W122 in Src SH3 and C100 in Abl SH3 were primarily responsible for loop conformation as we do not see specific W122 interactions with the side chain atoms of N116, and because these residues are within the core of the domain and positioned to regulate spacing/orientation of the loop.

**Figure 5.**
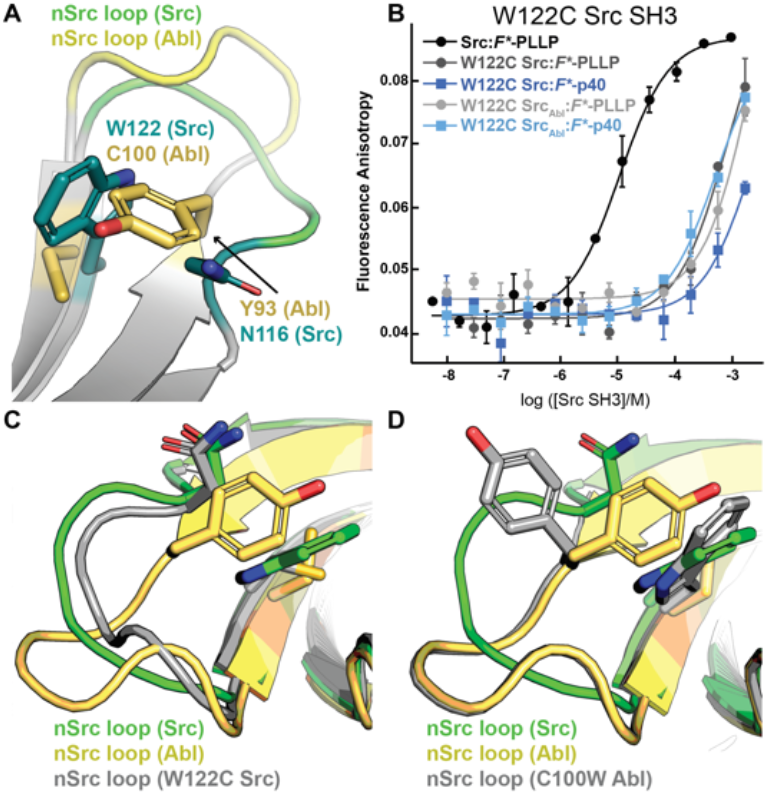
The WX motif following the nSrc loop determines its conformation. All structures, generated by Alphafold – Multimer, are shown in cartoon representation and colored as labeled. Side chain sticks are shown for relevant positions, and colored by atom (C=yellow (Abl), green (Src), or gray (W122C Src or C100W Abl), O=red, N=blue). (**A**) The X position of the WX motif in Src (X=W) and Abl (X=C) makes intra-SH3 interactions with N116 (Src) or Y93 (Abl). (**B**) Binding affinity data by fluorescence anisotropy experiments in triplicate confirms an importance for the W122 residue in the Src SH3 domain (**Table 1**). All averaged curves were normalized to an identical starting value to better illustrate Kd shifts. (**C-D**) The nSrc loop in the W122C Src model is shifted with respect to wild-type Src SH3 (**C**); whereas the nSrc loop in the C100W Abl SH3 model maintains an “Abl-like” conformation (**D**).

To test this hypothesis, we recombinantly expressed and purified a W122C mutation in Src SH3 and Src_Abl_ SH3, as described in the Materials and Methods. While W122C Src SH3 showed little to no binding for either peptide, the W122C Src_Abl_ SH3 domain was able to bind *F\**-p40 weakly, *K*D = 450 ± 190 μM (**Figure 5B**, **Table 1**). We predict this result may be due to a elevated “Abl-like” orientation of p40 binding to the Src_Abl_ SH3 domain, resulting in a weaker effect based on nSrc loop conformation (**Figure 4D**).

We also recombinantly expressed and purified C100W Abl SH3, as described in the Materials and Methods. The C100W Abl_Src_ SH3 protein was unstable in our purification. We saw that C100W Abl SH3 revealed undetectable binding to *F\**-PLLP, consistent with our observation that this peptide requires the lower “Src- like” peptide binding orientation due to the P4 Arg residue in *F\**-PLLP, which is likely not accessible in this mutant. Binding of C100W Abl SH3 to *F*-*p40 was weaker by ∼18-fold as compared to the wild-type protein, *K*D = 55 ± 20 μM, but was still relatively high for the SH3 domain interactions tested (**Figure 5B**, **Table 1**). Again, this is consistent with the elevated “Abl-like” peptide-binding orientation based on the Abl SH3 RT loop sequence.

Indeed, protein structure prediction models generated using Alphafold – Multimer show differences for nSrc loop conformation in the W122C Src SH3 and C100W Abl SH3 mutations. The W122C mutation in Src SH3 allows the nSrc loop to adopt a more “Abl-like” conformation at its C-terminus, although it is still mostly “Src-like” (**Figure 5C**). In contrast, we see no difference in the nSrc loop for a C100W Abl SH3 model as compared to the wild-type protein (**Figure 5D**). This is consistent with our biochemical assay results.

### Investigating a loop orientation-determining residue in the WX sequence motif

We next wanted to see how this position broadly affects loop orientation. We analyzed 39 previously solved SH3 structures, most of which had peptides bound in either the Class I (4 structures) or Class II (35 structures) orientation (*57*). We aligned these structures and colored the nSrc loop residues based on the “WX” motif immediately following the nSrc loop, including blue(s) for X = aromatic and yellow/orange for X = Cys or Leu (**Figure 6A**). Here, the “X” residues are the W122 Src SH3 and C100 Abl SH3 positions previously discussed. Although there are multiple structures for several of the SH3 domains, we have representative structures included for SH3 domains from the following proteins: WC (Abl), WL (CMS N-terminal SH3, or CMS-N, p40^phox^, p67^phox^), WY (CSK, Grb2-N, IB1), and WW (μ-PIX, CIN85-N, Cortactin, Fyn, GADS C-terminal SH3 or GADS-C, Hck, Itk, p47^phox^, PLC-γ1, SLA1, Src, STAM2) (*9*, *11*, *17*, *19*, *26–56*).

**Figure 6.**
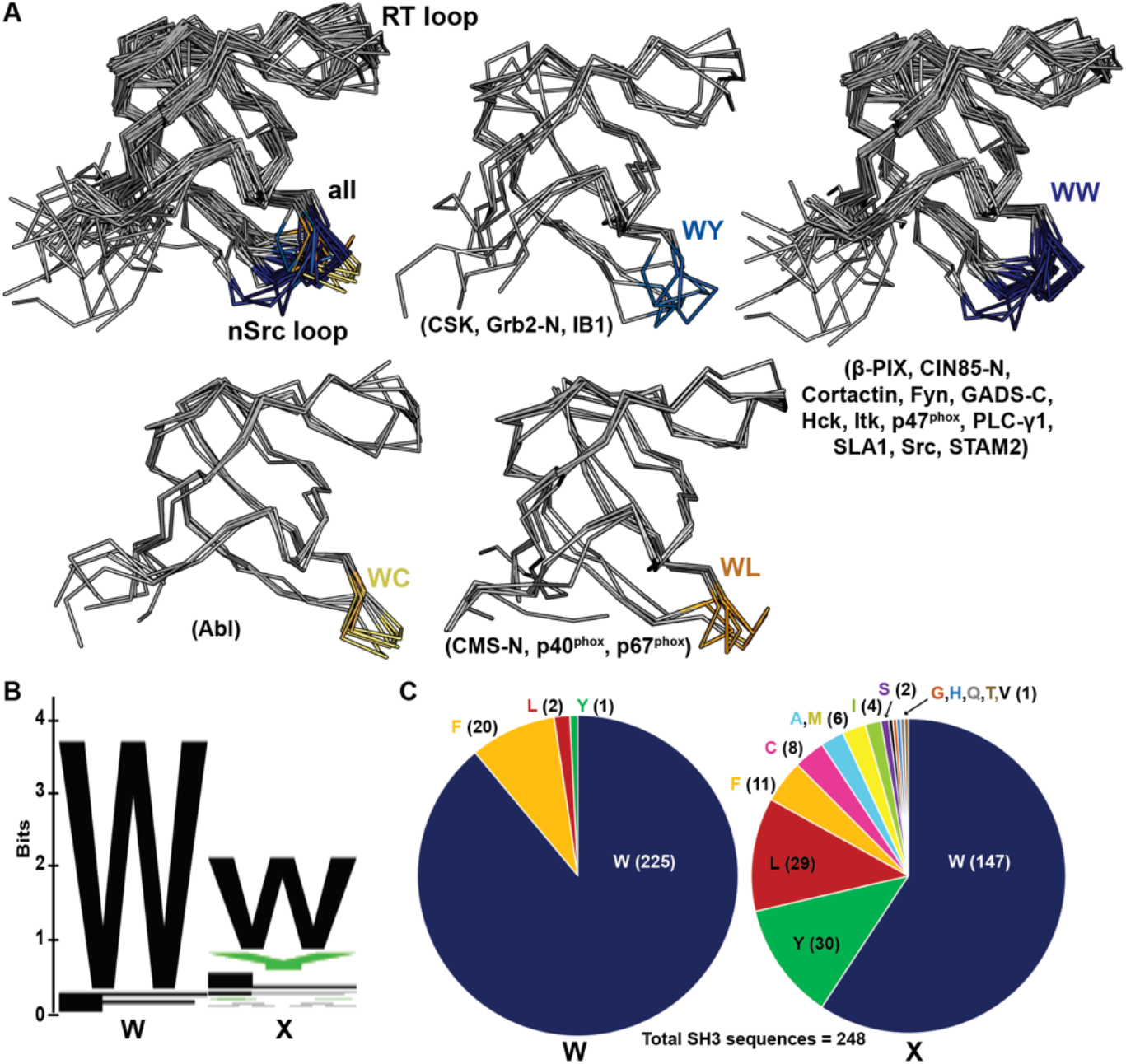
nSrc loop conformations and sequences of multiple SH3 domain structures. (**A**) All structures are shown in ribbon representation, with the nSrc loop residues colored based on the “WX” motif immediately following the loop (WW = dark blue, WY = blue, WC = yellow, WL = orange). The SH3 domains that are represented are indicated in parentheses; the majority of these structures include ligand, most in the Class II orientation. PDB ID codes used include: 1BBZ, 1JU5, 1AB0, 1OPL, 2FO0, 1OPK, 2J6F, 2J6O, 1K4U, 1W70, 1AWJ, 1YWO, 1RLP, 1H3H, 1UTI, 1UJ0, 1ZSG, 2P4R, 1CKA, 2DF6, 1AVZ, 1EFN, 1OEB, 2D0N, 2JT4, 2D1X, 2AK5, 2BZ8, 1WLP, 2HCK, 1AD5, 1FMK, 1Y57, 2SRC, 1KSW, 1JEG, 1AZE, 4GBQ, 2FPD (9, 11, 17, 19, 26–56). (**B-C**) The same data shown as a (**B**) WebLogo of the WX motifs for 253 SH3 domain sequences from UniProt, and (**C**) pie charts of the sequence composition per position. Of the 248 sequences analyzed, 91% contain a Trp in the first position and 59% in the second position.

Our analyses revealed that SH3 domains with WW or WY motifs following the nSrc loop contain more “Src-like” conformations whereas those with WC or WL motifs are “Abl-like.” Of the structures analyzed, the WY motif-containing SH3 domains show the most variability (**Figure 6A**). Taken together, our results suggest that this position plays an important role in the conformation of the nSrc loop. In combination with RT loop residues, this directly determines how ligands interact with the peptide-binding cleft and previously identified specificity pockets for SH3 domains.

Finally, we analyzed the sequences of all 344 annotated SH3 domains in the human proteome, using the UniProt database (*58–60*). We extracted the SH3 domain sequences, performed a multiple sequence alignment using Clustal Omega, and visualized the results using Jalview (**Figure S1**) (*61*, *62*). There were several atypical SH3 domains based on secondary structural elements according to manual visualization of available Alphafold structures (*63*, *64*); therefore, we chose to exclude any sequences that did not align properly from further analyses. In total, we analyzed the WX sequence motifs for 248 human SH3 domains (**Figures 6B-C**), and found that indeed, a WW sequence is the most observed, with 138 occurrences (55.6% of total sequences). Position by position, we see a Trp in the first position of the sequence motif in 91% of SH3 domains analyzed, and 59% in the second position (**Figure 6C**). While there were four alternative residues in the first position (including Leu, Phe, and Tyr), there are an additional thirteen alternatives in the second position (Ala, Cys, Phe, Gly, His, Ile, Leu, Met, Gln, Ser, Thr, Val, Tyr) (**Figure 6C**). Overall, this result confirms variability in the X, or second, position of the WX sequence motif defined here, which plays a role in SH3 specificity.

## Discussion

Despite their relatively small size, SH3 domains display remarkable plasticity in ligand binding and the family has been described as *versatile* and *diverse* (*4*, *13*, *57*, *65*). This includes the ability to bind ligands in opposite orientations, depending on the sequences of the short linear motifs (SLiMs) recognized. Specificity-determining components of SH3 have been described, and the RT and nSrc loops are known to play an important role (*13*, *57*, *65*). A conserved Trp residue immediately following the nSrc loop, and at the start of the βC strand was also identified as forming a hydrophobic pocket that is critical for ligand binding (*65*). Here, we extend discussion of this Trp residue to include its C-terminal neighbor, and define the “WX” motif, where the identity of the X residue is also important for specificity.

Calculated binding affinities and Alphafold – Multimer modeling of the Src and Abl SH3 domains, as well as chimeric proteins with the RT and nSrc loops switched in each, indicated that the conformation of these loops retains its wild-type position even in our chimeras. Our data suggests that specific amino acids in these loops determines how Class I peptides bind in these pockets, with “elevated Abl-like” and “lowered Src-like” orientations, the ability to access which directly determines relative binding affinities. The WX motif dictates the conformation of the nSrc loop and thus, affects ligand orientation. When we analyzed 39 SH3 domain structures, we saw clear patterns in the positions of the nSrc loops (**Figure 6**), supporting our biochemical and modeling data. Furthermore, the WX sequence motifs of 253 human SH3 domains confirmed that the X position is variable, with a total of fourteen amino acids in these sequences.

While this study was relatively limited in scope, our work identifies an important residue for loop positioning, and therefore SH3 specificity. It would be interesting to further test these observations using additional SH3 domains and a variety of ligand sequences. Future work could also include broad mutational characterization and analyses of the X position in the WX sequence motif identified, as well as high throughput studies of the specificity profiles of these differing SH3 domains. Considering their widespread and critical roles in the cell, a better understanding of the molecular determinants of SH3 specificity can advance knowledge of complex signaling pathways towards the improvement of human health.

## Materials and Methods

### Protein expression and purification

Sequences corresponding to the Abl and Src SH3 (residues 84-145, UniProt ID SRC_HUMAN) domains were used for the wild-type SH3 domains. All proteins were expressed as His6-SUMO fusion proteins using the pET28a(+) plasmid (Genscript). Expression and purification protocols were similar to those used previously for other SLiM-binding domains (*66–68*). All sequences used are in the **Supporting Information**.

Briefly, plasmids were transformed into BL21 DE3 chemically competent *Escherichia coli* cells. Selected colonies were grown in Terrific Broth (TB) at 37°C with shaking at 210 rpm. Once an optical density (OD) of 0.6-0.8 at λ = 600 nm was reached, protein expression was induced by the addition of 0.15 mM isopropyl-β-D-1-thiogalactopyranoside (IPTG) for 16-18 h at 18°C. Cells were harvested via centrifugation at 3,000x*g* for 10 min, followed by resuspension in lysis buffer (0.05 M Tris pH 7.5, 0.2 M NaCl, 0.01 M MgCl2, 0.01 M CaCl2, 0.05 M imidazole pH 7.5, 20% (*w/v*) glycerol, 0.25 mM TCEP; 20 μg/mL DNAse and Complete EDTA-free protease inhibitor tablets (1 tablet/50 mL lysis buffer) were also added). Cells were lysed using sonication at 4° C and whole cell lysate was clarified with centrifugation at 17,500 rpm at 4° C for 30 min. The filtered supernatant was applied to a NiNTA HisTrap 5 mL column (Cytiva) with wash buffer (0.025 M Tris pH 7.5, 0.25 M NaCl, 10% (*w/v*) glycerol, 0.025 M imidazole pH 7.5, 0.25 mM tris(2- carboxyethyl)phosphine (TCEP)). Protein was eluted into fractions using elution buffer (0.025 M Tris-HCl pH 7.5, 0.05 M NaCl, 10%(*w/v*) glycerol, 0.4 M imidazole pH 7.5, 0.25mM TCEP). Pooled fractions were dialyzed in the presence of ULP-1 SUMO protease in dialysis buffer (0.025 M Tris pH 7.5, 0.15 M NaCl, 10% (*w/v*) glycerol, 0.5mM TCEP) overnight, and then run over another NiNTA His-Trap 5 mL column to separate the SH3 domain from the His6-SUMO tag, using running buffer plus 0.025 M imidazole pH 7.5. The flow-through was concentrated using Amicon Ultra 3K Centrifugal Filters and further purified using a Superdex S75 16/600 column, with running buffer. SDS-PAGE was used to assess protein purity and the absorbance at λ = 280 nm with the calculated extinction coefficient(s), by Expasy ProtParam, was used to calculate protein concentration (*68*).

Calculated extinction coefficients used were as follows: SUMO-Src-SH3 = 18,450 M^-1^ cm^-1^ (cleaved, 16,960 M^-1^ cm^-1^); SUMO-Abl-SH3 = 16,960 M^-1^ cm^-1^ (cleaved, 15,470 M^-1^ cm^-1^); SUMO-Src_Abl_- SH3 = 18,450 M^-1^ cm^-1^ (cleaved, 16,960 M^-1^ cm^-1^); SUMO-Abl_Src_-SH3 = 16,960 M^-1^ cm^-1^ (cleaved, 15,470 M^-1^ cm^-1^); W122C SUMO-Src-SH3 = 12,950 M^-1^ cm^-1^ (cleaved, 16,460 M^-1^ cm^-1^); C100W SUMO-Abl-SH3 = 22,460 M^-1^ cm^-1^ (cleaved, 20,970 M^-1^ cm^-1^); W122C SUMO-Src_Abl_-SH3 = 12,950 M^-1^ cm^-1^ (cleaved, 11,460 M^-1^ cm^-1^).

### Fluorescence anisotropy assays

*The* peptides used for fluorescence anisotropy assays were *F*\*-PLLP (sequence: FITC-Ahx-LASRPLPLLP), the Src SH3-specific binder, and *F*\*-p40 (FITC-Ahx-APTYSPPPPP), the Abl SH3-specific binder (Biomatik) (*6*, *20*). Fluorescence anisotropy assays were performed using a BioTek Synergy H1 plate reader, as previously described (*66–69*). Solution conditions included running buffer plus 0.1 mg/mL bovine serine albumin (BSA) and 0.5 mM Thesit, with 30 nM fluorescent peptide. Determined *K*D values were the average calculated from triplicate experiments (*70*).

### Programs used for structural modeling and analyses

AlphaFold – Multimer (in the CoLabFold notebook) was used to model ligand-bound SH3 domains (*23*, *24*). Binding curves were visualized using KaleidaGraph (Synergy software). Structural figures were rendered using PyMOL. We used UniProt to download the human SH3ome, following by Clustal Omega to generate a multiple sequence alignment, Jalview to view the results, and WebLogo and Excel to analyze and visualize the data (**Figure S1**) (*58–62*, *71*). In our initial curation of the human SH3 multiple sequence alignment, we used the Alphafold protein database to manually visualize WX sequence motifs that did not properly align (*63*, *64*).

## Contributions (CRediT Classification)

**Frederick Longshore-Neate**: conceptualization (equal), data curation (equal), formal analysis (equal), funding acquisition (supporting), investigation (lead), validation (supporting), visualization (equal), writing – original draft preparation (supporting). **Caroline Ceravolo**: formal analysis (equal), funding acquisition (supporting), investigation (equal), writing – review & editing (supporting). **Cole Masuga**: investigation (supporting). **Elise F. Tahti**: investigation (supporting). **Jadon M. Blount**: investigation (supporting). **Sarah N. Smith**: investigation (supporting). **Jeanine F. Amacher**: conceptualization (equal), data curation (equal), formal analysis (equal), funding acquisition (lead), project administration (lead), resources (lead), supervision (lead), visualization (equal), writing – original draft preparation (lead).

## Acknowledgements

The authors would like to thank all members of the Amacher lab for thoughtful discussions and general research support. We would also like to thank Dr. Lionel (Lee) Brooks, who was instrumental in providing computational support towards downloading the human SH3 domain sequences.

