## Supporting Information for "The conformation of the nSrc specificity-determining loops in the Src and Abl SH3 domains are modulated by a “WX” conserved sequence motif"

Table of Contents:

**Figure S1. Alignment of human SH3 domains.** 2

**Protein sequences used.** 6

**Figure S1. Alignment of 248 human SH3 domains extracted from the UniProt database.** Where sequences were variable, e.g., differing lengths of the RT and nSrc loops, the alignment was truncated for clarify. Asterisks (*) are used to show where the alignment was edited. Asterisks (*) on the left also highlight the Abl (UniProt ID P00519) and Src (UniProt ID P12931) sequences.

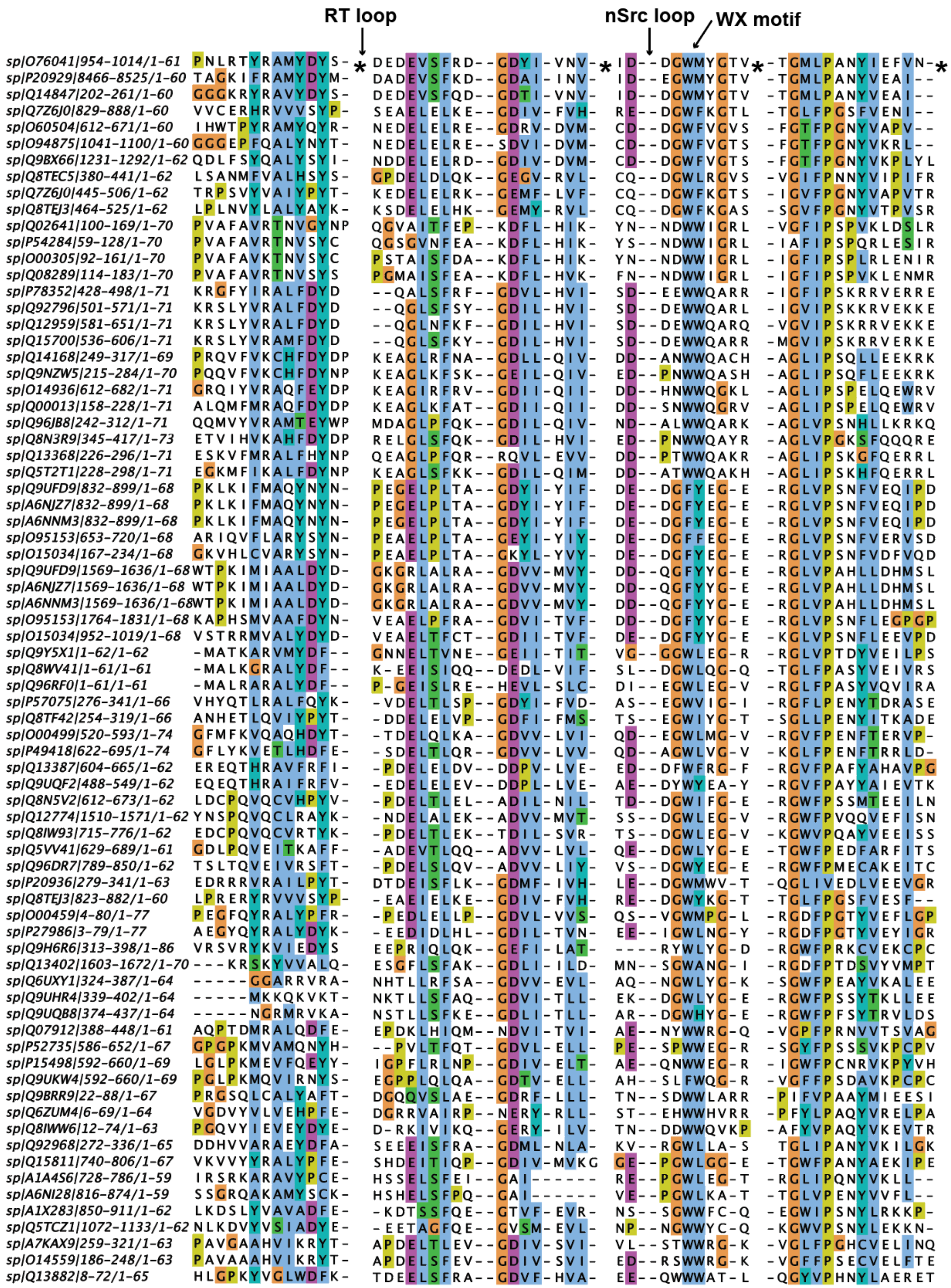

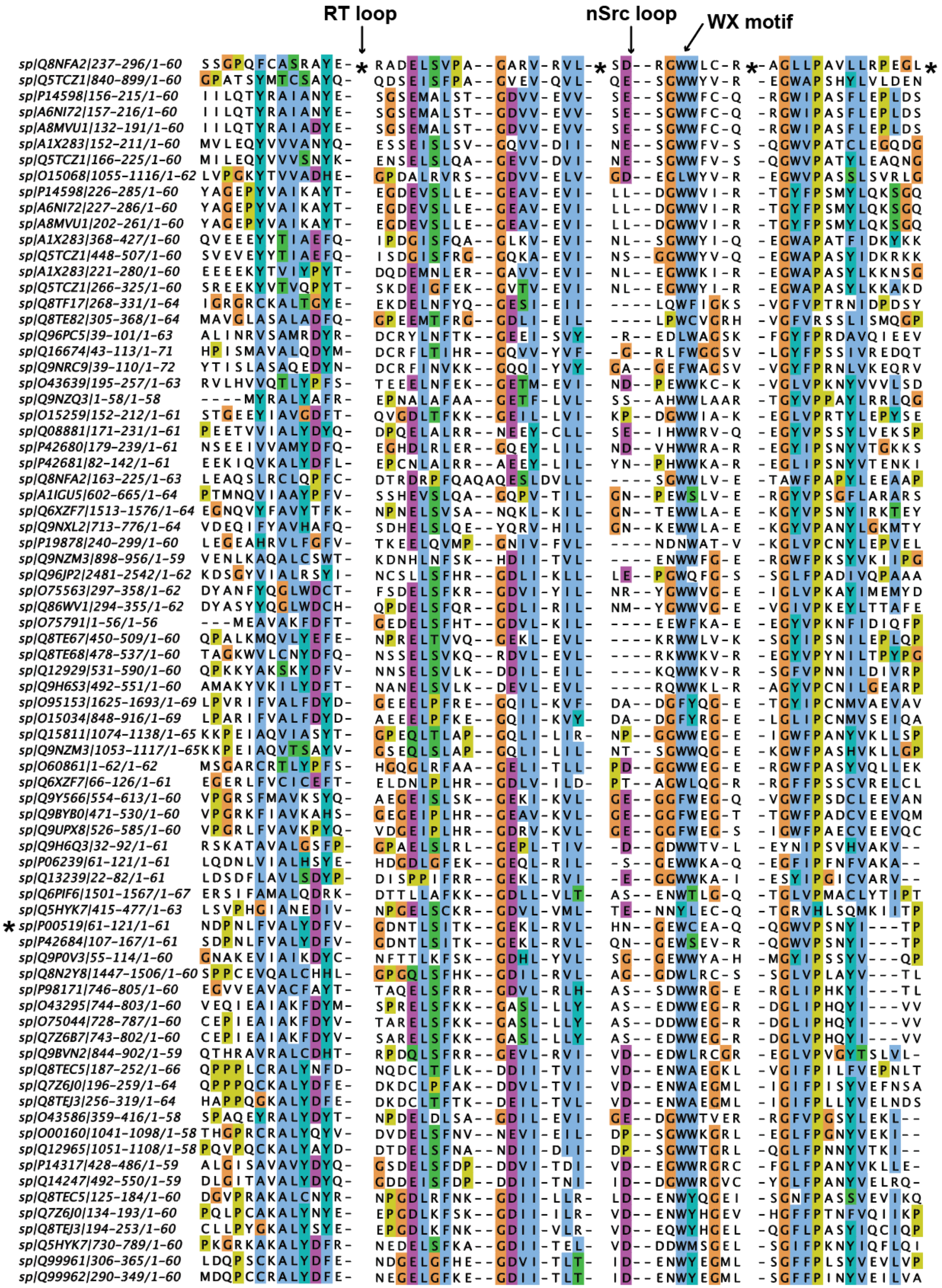
RT loop
nSrc loop
WX motif

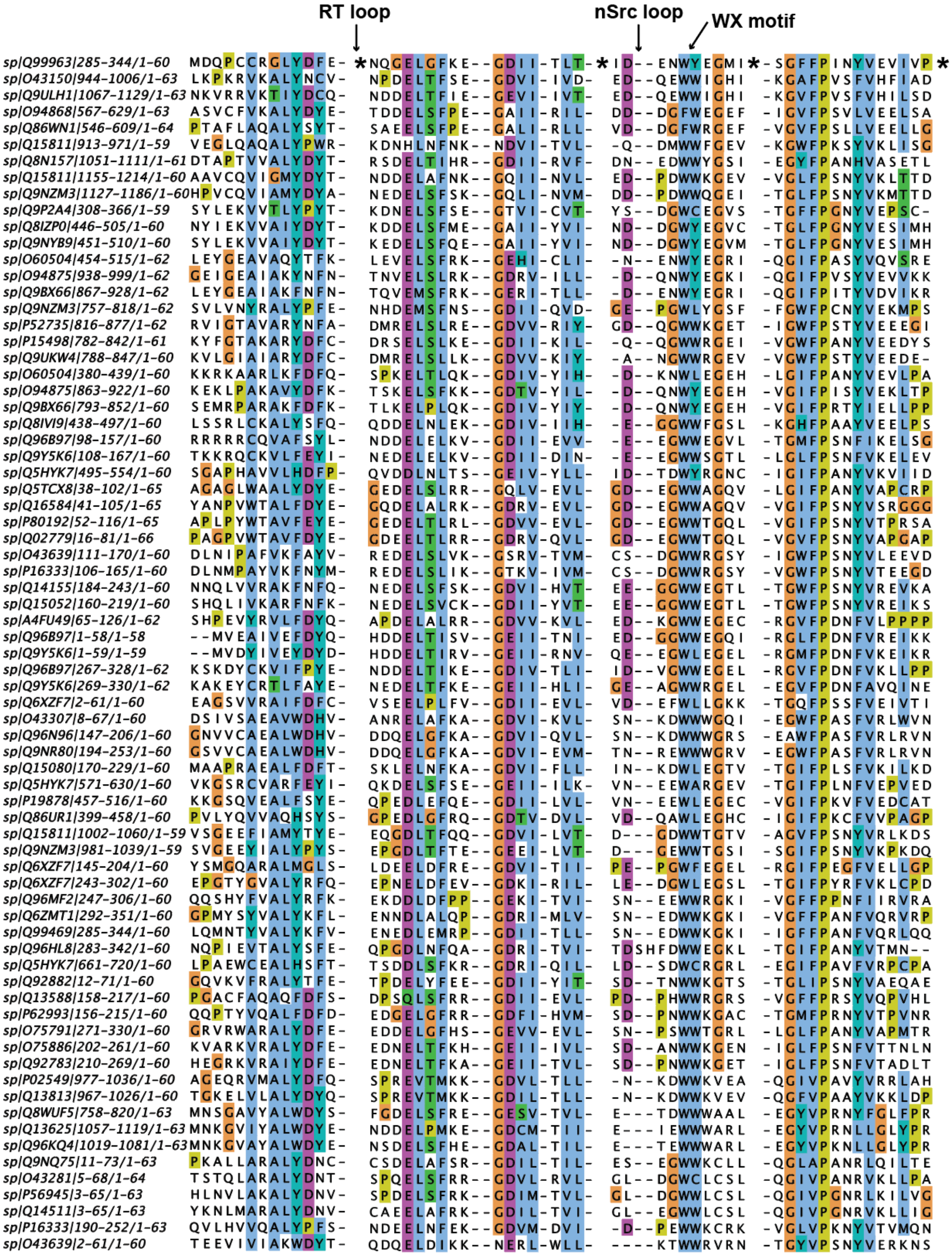
RT loop
nSrc loop
WX motif
sp|Q99963|285-344/1-60 MDQPCCRGLYDFE- * NQGELGFKE--GDII-TLT- * ID--ENWYEGMI * -SGFFPINYVEVIVP *
sp|O43150|944-1006/1-63 LKPKRVKALYNCV- NPDELTFSE--GDVI-IVD- ED--QEWWIGHI -KGAFPVSFVHFIAD
sp|Q9ULH1|1067-1129/1-63 NKVRRVKTIYDCQ- NDDELTFIE--GEVI-IVT- ED--QEWWIGHI -KGVFPVSFVHILSD
sp|O94868|567-629/1-63 ASVCFVKALYDYE- TDDELSFPE--GAII-RIL- DD--DGFWEGEF -IGVFPSVLVEELSA
sp|Q86WN1|546-609/1-64 PTAFLAQALYSYT- SAEELSFPE--GALI-RLL- VD--DGFWRGEF -VGVFPSLLVEELLG
sp|Q15811|913-971/1-59 VEGLQAQALYPWR- KDNHLNFNK--NDVI-TVL- -Q--DMWWFGEV -KGWFPKSYVKLISG
sp|Q8N157|1051-1111/1-61 DTAPTVVALYDYT- RSDELTIHR--GDII-RVF- DN--EDWWYGSI -EGYFPANHVASETL
sp|Q15811|1155-1214/1-60 AAVCQVIGMYDYT- NDDELAFNK--GQII-NVL- ED--PDWWKGEV -VGLFPSNYVKLTTD
sp|Q9NZM3|1127-1186/1-60 HPVCQVIAMYDYA- NEDELSFSK--GQLI-NVM- DD--PDWWQGEI -TGLFPSNYVKMTTD
sp|Q9P2A4|308-366/1-59 SYLEKVVTLYPYT- KDNELSFSE--GTVI-CVT- YS--DGWCEGVS -TGFFPGNYVEPSC-
sp|Q8IZP0|446-505/1-60 NYIEKVVAIYDYT- KDDELSFME--GAII-YVI- ND--DGWYEGVC -TGLFPGNYVESIMH
sp|Q9NYB9|451-510/1-60 SYLEKVVAIYDYT- KEDELSFQE--GAII-YVI- ND--DGWYEGVM -TGLFPGNYVESIMH
sp|O60504|454-515/1-62 LEYGEAVAQYTFK- LEVELSFRK--GEHI-CLI- -N--ENWYEGRI -QGIFPASYVQVSRE
sp|O94875|938-999/1-62 GEIGEAIAKYNFN- TNVELSLRK--GDRV-ILL- -D--QNWYEGKI -QGIFPVSYVEVVKK
sp|Q9BX66|867-928/1-62 LEYGEAIAKFNFN- TQVEMSFRK--GERI-TLL- -D--ENWYEGRI -QGIFPITYVDVIKR
sp|Q9NZM3|757-818/1-62 SVLVNYRALYPFE- NHDEMSFNS--GDII-QVD- GE--PGWLYGSF -FGWFPCNYVEKMPS
sp|P52735|816-877/1-62 RVIGTAVARYNFA- DMRELSLRE--GDVV-RIY- GD--QGWWKGET -IGWFPSTYVEEEGI
sp|P15498|782-842/1-61 KYFGTAKARYDFC- DRSELSLKE--GDII-KIL- -Q--QGWWRGEI -VGWFPANYVEDYS
sp|Q9UKW4|788-847/1-60 KVLGIAIARYDFC- DMRELSLLK--GDVV-KIY- -A--NGWWRGEV -VGWFPSTYVEEDE-
sp|O60504|380-439/1-60 KKRKAARLKFDFQ- SPKELTLQK--GDIV-YIH- -D--KNWLEGEH -LGIFPANYVEVLPA
sp|O94875|863-922/1-60 KEKLPAKAVYDFK- TSKELSFKK--GDTV-YIL- -D--QNWYEGEH -VGIFPISYVEKLTP
sp|Q9BX66|793-852/1-60 SEMRPARAKFDFK- TLKELPLQK--GDIV-YIY- -D--QNWYEGEH -VGIFPRTYIELLPP
sp|Q8IVI9|438-497/1-60 LSSRLCKALYSFQ- QDDELNLEK--GDIV-IIH- -E--GGWWFGSL -KGHFPAAYVEELPS
sp|Q96B97|98-157/1-60 RRRRRCQVAFSYL- NDDELELKV--GDII-EVV- -E--EGWWEGVL -TGMFPSNFIKELSG
sp|Q9Y5K6|108-167/1-60 TKKRQCKVLFEYI- NEDELELKV--GDII-DIN- -E--EGWWSGTL -LGLFPSNFVKELEV
sp|Q5HYK7|495-554/1-60 SGAPHAVVLHDFP- QVDDLNLTS--GEIV-YLL- ID--TDWYRGNC -IGIFPANYVKVIID
sp|Q5TCX8|38-102/1-65 AGAGLWAALYDYE- GEDELSLRR--GQLV-EVL- GD--EGWWAGQV -LGIFPANYVAPCRP
sp|Q16584|41-105/1-65 YANPVWTALFDYE- GQDELALRK--GDRV-EVL- GD--EGWWAGQV -VGIFPSNYVSRGGG
sp|P80192|52-116/1-65 APLPYWTAVFEYE- GEDELTLRL--GDVV-EVL- GD--EGWWTGQL -VGIFPSNYVTPRSA
sp|Q02779|16-81/1-66 PAGPVWTAVFDYE- GDEELTLRR--GDRV-QVL- GD--EGWWTGQL -VGVFPSNYVAPGAP
sp|O43639|111-170/1-60 DLNIPAFVKFAYV- REDELSLVK--GSRV-TVM- CS--DGWWRGSY -IGWFPSNYVLEEVD
sp|P16333|106-165/1-60 DLNMPAYVKFNYM- REDELSLIK--GTKV-IVM- CS--DGWWRGSY -VGWFPSNYVTEEGD
sp|Q14155|184-243/1-60 NNQLVVRAKFNFQ- NEDELSFSK--GDVI-HVT- EE--GGWWEGTL -TGWFPSNYVREVKA
sp|Q15052|160-219/1-60 SHQLIVKARFNFK- NEDELSVCK--GDII-YVT- EE--GGWWEGTL -TGWFPSNYVREIKS
sp|A4FU49|65-126/1-62 SHPEVYRVLFDYQ- APDELALRR--GDVV-KVL- ED--KGWWEGEC -RGVFPDNFVLPPPP
sp|Q96B97|1-58/1-58 --MVEAIVEFDYQ- HDDELTISV--GEII-TNI- ED--GGWWEGQI -RGLFPDNFVREIKK
sp|Q9Y5K6|1-59/1-59 --MVDYIVEYDYD- HDDELTIRV--GEII-RNV- QE--EGWLEGEL -RGMFPDNFVKEIKR
sp|Q96B97|267-328/1-62 KSKDYCKVIFPYE- NDDELTIKE--GDIV-TLI- ID--VGWWEGEL -RGVFPDNFVKLLPP
sp|Q9Y5K6|269-330/1-62 KAKEYCRTLFAYE- NEDELTFKE--GEII-HLI- GE--AGWWRGEL -EGVFPDNFAVQINE
sp|Q6XZF7|2-61/1-60 EAGSVVRAIFDFC- VSEELPLFV--GDII-EVL- VD--EFWLLGKK -TGQFPSSFVEIVTI
sp|O43307|8-67/1-60 DSIVSAEAVWDHV- ANRELAFKA--GDVI-KVL- SN--KDWWWGQI -EGWFPASFVRLWVN
sp|Q96N96|147-206/1-60 GNVVCAEALWDHV- DDQELGFKA--GDVI-QVL- SN--KDWWWGRS -EAWFPASFVRLRVN
sp|Q9NR80|194-253/1-60 GSVVCAEALWDHV- DDQELGFKA--GDVI-EVM- TN--REWWWGRV -EGWFPASFVRLRVN
sp|Q15080|170-229/1-60 MAAPRAEALFDFT- SKLELNFKA--GDVI-FLL- IN--KDWLEGTV -TGIFPLSFVKILKD
sp|Q5HYK7|571-630/1-60 VKGSRCVARFEYI- QKDELSFSE--GEII-ILK- VN--EEWARGEV -TGIFPLNFVEPVED
sp|P19878|457-516/1-60 KKGSQVEALFSYE- QPEDLEFQE--GDII-LVL- VN--EEWLEGEC -VGIFPKVFVEDCAT
sp|Q86UR1|399-458/1-60 PVLYQVVAQHSYS- GPEDLGFRQ--GDTV-DVL- VD--QAWLEGHC -IGIFPKCFVVPAGP
sp|Q15811|1002-1060/1-59 VSGEEFIAMYTYE- EQGDLTFQQ--GDVI-LVT- D---GDWWTGTV -AGVFPSNYVRLKDS
sp|Q9NZM3|981-1039/1-59 SVGEEYIALYPYS- EPGDLTFTE--GEEI-LVT- D---GEWWTGSI -SGIFPSNYVKPKDQ
sp|Q6XZF7|145-204/1-60 YSMGQARALMGLS- LDEELDFRE--GDVI-TII- PE--PGWFEGEL -RGIFPEGFVELLGP
sp|Q6XZF7|243-302/1-60 EPGTYGVALYRFQ- EPNELDFEV--GDKI-RIL- LE--DGWLEGSL -TGIFPYRFVKLCPD
sp|Q96MF2|247-306/1-60 QQSHYFVALYRFK- EKDDLDFPP--GEKI-TVI- SN--EEWWRGKI -VGFFPPNFIIRVRA
sp|Q6ZMT1|292-351/1-60 GPMYSYVALYKFL- ENNDLALQP--GDRI-MLV- SN--EDWWKGKI -VGFFPANFVQRVRP
sp|Q99469|285-344/1-60 LQMNTYVALYKFV- ENEDLEMRP--GDII-TLL- SN--EDWWKGKI -IGFFPANFVQRLQQ
sp|Q96HL8|283-342/1-60 NQPIEVTALYSFE- QPGDLNFQA--GDRI-TVI- TDSHFDWWEGKL -TGIFPANYVTMN--
sp|Q5HYK7|661-720/1-60 LPAEWCEALHSFT- TSDDLSFKR--GDRI-QIL- LD--SDWCRGRL -EGIFPAVFVRPCPA
sp|Q92882|12-71/1-60 GQVKVFRALYTFE- TPDELYFEE--GDII-YIT- SD--TNWWKGTS -TGLIPSNYVAEQAE
sp|Q13588|158-217/1-60 PGACFAQAQFDFS- DPSQLSFRR--GDII-EVL- PD--PHWWRGRS -VGFFPRSYVQPVHL
sp|P62993|156-215/1-60 QQPTYVQALFDFD- EDGELGFRR--GDFI-HVM- SD--PNWWKGAC -TGMFPRNYVTPVNR
sp|O75791|271-330/1-60 GRVRWARALYDFE- EDDELGFHS--GDVV-EVL- SN--PSWWTGRL -LGLFPANYVAPMTR
sp|O75886|202-261/1-60 KVARKVRALYDFE- EDNELTFKH--GEII-IVL- SD--ANWWKGEN -IGLFPSNFVTTNLN
sp|Q92783|210-269/1-60 HEGRKVRAIYDFE- EDNELTFKA--GEII-TVL- SD--PNWWKGET -IGLFPSNFVTADLT
sp|P02549|977-1036/1-60 AGEQRVMALYDFQ- SPREVTMKK--GDVL-TLL- -N--KDWWKVEA -QGIVPAVYVRRLAH
sp|Q13813|967-1026/1-60 TGKELVLALYDYQ- SPREVTMKK--GDIL-TLL- -N--KDWWKVEV -QGFVPAAYVKKLDP
sp|Q8WUF5|758-820/1-63 MNSGAVYALWDYS- FGDELSFRE--GESV-TVL- E---TDWWWAAL -EGYVPRNYFGLFPR
sp|Q13625|1057-1119/1-63 MNKGVIYALWDYE- NDDELPMKE--GDCM-TII- E---IEWWWARL -EGYVPRNLLGLYPR
sp|Q96KQ4|1019-1081/1-63 MNKGVAYALWDYE- NSDELSFHE--GDAL-TIL- E---TEWWWARL -EGYVPKNLLGLYPR
sp|Q9NQ75|11-73/1-63 PKALLARALYDNC- CSDELAFSR--GDIL-TIL- ES--EGWWKCLL -QGLAPANRLQILTE
sp|O43281|5-68/1-64 TSTQLARALYDNT- SPQELSFRR--GDVL-RVL- GL--DGWCLCSL -QGIVPANRVKLLPA
sp|P56945|3-65/1-63 HLNVLAKALYDNV- SPDELSFRK--GDIM-TVL- GL--DGWWLCSL -QGIVPGNRLKILVG
sp|Q14511|3-65/1-63 YKNLMARALYDNV- CAEELAFRK--GDIL-TVI- GL--EGWWLCSL -QGIVPGNRVKLLIG
sp|P16333|190-252/1-63 QVLHVVQALYPFS- NDEELNFEK--GDVM-DVI- -D--PEWWKCRK -VGLVPKNYVTVMQN
sp|O43639|2-61/1-60 TEEVIVIAKWDYT- QDQELDIKK--NERL-WLL- ----KTWWRVRN -TGYVPSNYVERKNS

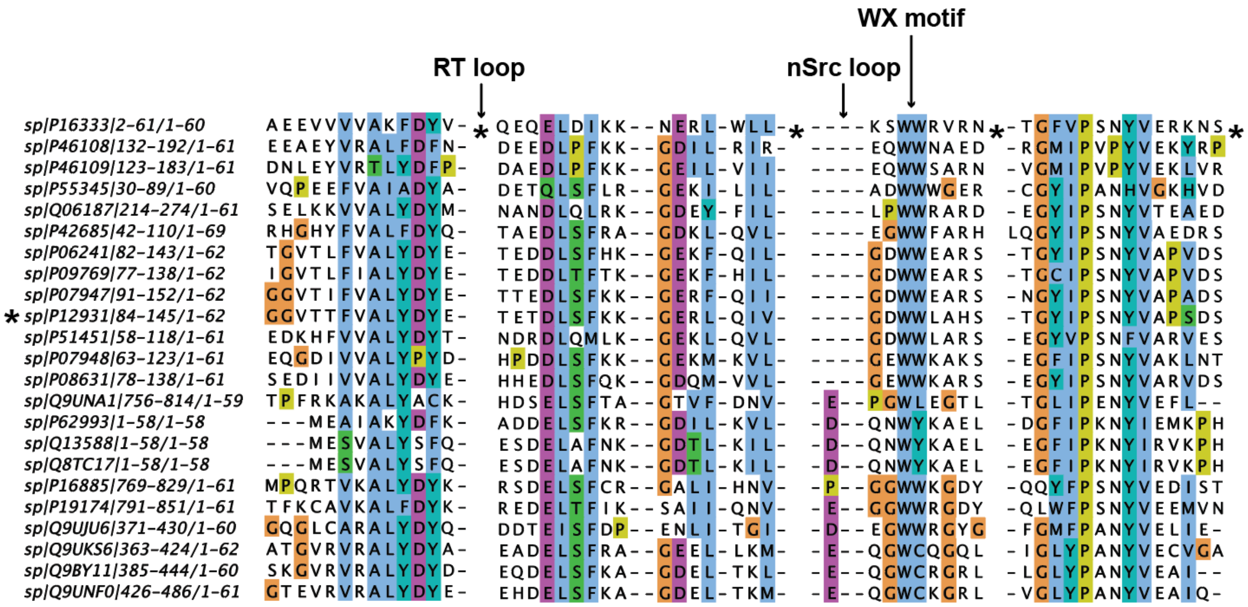
WX motif
RT loop
nSrc loop
sp|P16333|2-61/1-60
sp|P46108|132-192/1-61
sp|P46109|123-183/1-61
sp|P55345|30-89/1-60
sp|Q06187|214-274/1-61
sp|P42685|42-110/1-69
sp|P06241|82-143/1-62
sp|P09769|77-138/1-62
sp|P07947|91-152/1-62
sp|P12931|84-145/1-62
sp|P51451|58-118/1-61
sp|P07948|63-123/1-61
sp|P08631|78-138/1-61
sp|Q9UNA1|756-814/1-59
sp|P62993|1-58/1-58
sp|Q13588|1-58/1-58
sp|Q8TC17|1-58/1-58
sp|P16885|769-829/1-61
sp|P19174|791-851/1-61
sp|Q9UJU6|371-430/1-60
sp|Q9UKS6|363-424/1-62
sp|Q9BY11|385-444/1-60
sp|Q9UNF0|426-486/1-61

**Recombinant protein sequences used in this study.** The pET28a(+) plasmid was used for all. The 6xHis-tag and TEV protease site included are in **bold** and the SUMO (*Saccharomyces cerevisiae* Smt3, NCBI reference: NP_010798.1) sequence is underlined. In chimeric constructs and mutants, the changed residue(s) is/are in **red**.

>SUMO_Src_SH3
**MGSSHHHHHHGSGLVPRGSAS**MSDSEVNQEAKPEVKPEVKPETHINLKVSDGSSEIFFKIKKTTPLRRLM
EAFAKRQGKEMDSLRFLYDGIRIQADQTPEDLDMEDNDIIEAHREQIGGVTTFVALYDYESRTETDLSFK
KGERLQIVNNTEGDWWLAHSLSTGQTGYIPSNYVAPSDS

>SUMO_Abl_SH3
**MGSSHHHHHHGSGLVPRGSAS**MSDSEVNQEAKPEVKPEVKPETHINLKVSDGSSEIFFKIKKTTPLRRLM
EAFAKRQGKEMDSLRFLYDGIRIQADQTPEDLDMEDNDIIEAHREQIGGNDPNLFVALYDFVASGDNTLS
ITKGEKLRVLGYNHNGEWCEAQTKNGQGWVPSNYITPVNS

>SUMO_Src$_{Abl}$_SH3
**MGSSHHHHHHGSGLVPRGSAS**MSDSEVNQEAKPEVKPEVKPETHINLKVSDGSSEIFFKIKKTTPLRRLM
EAFAKRQGKEMDSLRFLYDGIRIQADQTPEDLDMEDNDIIEAHREQIGGVTTFVALYDY**VASGDN**DLSFK
KGERLQIVNN**HNGE**WWLAHSLSTGQTGYIPSNYVAPSDS

>SUMO_Abl$_{Src}$_SH3
**MGSSHHHHHHGSGLVPRGSAS**MSDSEVNQEAKPEVKPEVKPETHINLKVSDGSSEIFFKIKKTTPLRRLM
EAFAKRQGKEMDSLRFLYDGIRIQADQTPEDLDMEDNDIIEAHREQIGGNDPNLFVALYDF**ESRTET**TLS
ITKGEKLRVLGYN**TEGD**WCEAQTKNGQGWVPSNYITPVNS

>SUMO_W122C_Src_SH3
**MGSSHHHHHHGSGLVPRGSAS**MSDSEVNQEAKPEVKPEVKPETHINLKVSDGSSEIFFKIKKTTPLRRLM
EAFAKRQGKEMDSLRFLYDGIRIQADQTPEDLDMEDNDIIEAHREQIGGVTTFVALYDYESRTETDLSFK
KGERLQIVNNTEGDW**C**LAHSLSTGQTGYIPSNYVAPSDS

>SUMO_W122C_Src$_{Abl}$_SH3
**MGSSHHHHHHGSGLVPRGSAS**MSDSEVNQEAKPEVKPEVKPETHINLKVSDGSSEIFFKIKKTTPLRRLM
EAFAKRQGKEMDSLRFLYDGIRIQADQTPEDLDMEDNDIIEAHREQIGGVTTFVALYDY**VASGDN**DLSFK
KGERLQIVNN**HNGE**W**C**LAHSLSTGQTGYIPSNYVAPSDS

>SUMO_C100W_Abl_SH3
**MGSSHHHHHHGSGLVPRGSAS**MSDSEVNQEAKPEVKPEVKPETHINLKVSDGSSEIFFKIKKTTPLRRLM
EAFAKRQGKEMDSLRFLYDGIRIQADQTPEDLDMEDNDIIEAHREQIGGNDPNLFVALYDFVASGDNTLS
ITKGEKLRVLGYNHNGEW**W**EAQTKNGQGWVPSNYITPVNS
